# Nuclease-dead S. *aureus* Cas9 downregulates alpha-synuclein and reduces mtDNA damage and oxidative stress levels in patient-derived stem cell model of Parkinson’s disease

**DOI:** 10.1101/2023.01.24.525105

**Authors:** Danuta Sastre, Faria Zafar, C. Alejandra Morato Torres, Desiree Piper, Deniz Kirik, Laurie H. Sanders, Stanley Qi, Birgitt Schüle

**Author notes:** corresponding author, **Corresponding author** Birgitt Schüle, MD, Dr. med., Stanford University School of Medicine, Department of Pathology, 300 Pasteur Dr., R271/217, Stanford, CA 94305. these authors contributed equally to this work (Danuta Sastre, Faria Zafar). SRI International, Biosciences Division, Menlo Park, 94025 CA, U.S.A.

## Abstract

Parkinson’s disease (PD) is one of the most common neurodegenerative diseases, but no disease modifying therapies have been successful in clinical translation presenting a major unmet medical need. A promising target is alpha-synuclein or its aggregated form, which accumulates in the brain of PD patients as Lewy bodies. While it is not entirely clear which alpha-synuclein protein species is disease relevant, mere overexpression of alpha-synuclein in hereditary forms leads to neurodegeneration.

To specifically address gene regulation of alpha-synuclein, we developed a CRISPR interference (CRISPRi) system based on the nuclease dead *S. aureus* Cas9 (SadCas9) fused with the transcriptional repressor domain Krueppel-associated box to controllably repress alpha-synuclein expression at the transcriptional level. We screened single guide (sg)RNAs across the *SNCA* promoter and identified several sgRNAs that mediate downregulation of alpha-synuclein at varying levels. CRISPRi downregulation of alpha-synuclein in iPSC-derived neuronal cultures from a patient with an *SNCA* genomic triplication showed functional recovery by reduction of oxidative stress and mitochondrial DNA damage.

Our results are proof-of-concept *in vitro* for precision medicine by targeting the *SNCA* gene promoter. The *SNCA* CRISPRi approach presents a new model to understand safe levels of alpha-synuclein downregulation and a novel therapeutic strategy for PD and related alpha-synucleinopathies.

## Introduction

Parkinson’s disease (PD) represents one of the two most common neurodegenerative diseases of aging, Alzheimer’s disease being the other. Approximately 1-2% of the population over 65 years of age are affected by this disorder, but the disease can also occur at an earlier age due to causative mutations and specific genetic susceptibility and/or exposure to environmental factors, drugs, or trauma^1-3^. While symptomatic forms of therapy are available, proven disease-modifying agents have yet to be discovered^4,5^.

In the last two decades, genetics has greatly advanced our understanding of PD and many newly discovered genes have become targets for therapeutic developments^6^. The first PD gene was alpha-synuclein (*SNCA*) in which causative mutations have been described that can be grouped into single point mutations and large copy number variants leading to genomic duplications and triplications on chromosome 4q22.1^7^. Clinically, there is well documented gene dosage effect with earlier onset and faster progression in cases with *SNCA* genomic triplications^8-10^. These medical genetic findings together with molecular and structural characterization of toxic alpha-synuclein aggregates and neuropathological evidence of alpha-synuclein accumulation in Lewy bodies has led to a wide-spread hypothesis that lowering alpha-synuclein levels could be of therapeutic benefit for PD and related alpha-synucleinopathies^4^. However, it is not clear what safe levels of alpha-synuclein downregulation are.

As a proof-of-concept that modulation of alpha-synuclein expression can lead to neuroprotection, the use of small interference RNAs (siRNA) in the brain has recently been shown be effective against endogenous murine alpha-synuclein ^11-14^. In a translational non-human primate model, siRNA showed consistent knockdown of alpha-synuclein in MPTP-exposed squirrel monkeys ^15^. Another line of evidence that lowering alpha-synuclein expression levels are neuroprotective shows that a small molecule beta2-adrenoreceptor agonist is a regulator of the *SNCA* gene by modifying the histone 3 lysine 27 acetylation of the *SNCA* promoter and ameliorates mitochondrial dysfunction in human stem cell-derived neuronal cultures with an *SNCA* genomic triplication^16^.

New versatile CRISPR gene-engineering technologies to precisely target positions in the human genome can also be exploited to lower alpha-synuclein mRNA expression level. Beyond editing, CRISPR/Cas can also activate or inhibit gene expression via steric hindrance of transcription itself or epigenetic changes such as DNA methylation or chromatin modification^17,18^. The first proof that nuclease-dead CRISPR/Cas9 regulates alpha-synuclein expression was published in 2016^19^. A limited number of small guide (sg)RNAs with a PAM recognition sequence for the 4.2kb *S. pyogenes* version of Cas9 were tested and one sgRNA was experimentally found to downregulate alpha-synuclein expression by approximately 50% in human cancer cell lines and human induced-pluripotent stem cell (iPSC) models^19^.

Here, we developed a different gene-engineering approach of a smaller nuclease-dead CRISPR/*S. aureus* Cas9 fused to Krueppel-associated box (KRAB) domains to regulate alpha-synuclein expression in a patient-derived iPSC model. In a screen of 32 sgRNAs, we identified a panel of sgRNAs that allows for regulating alpha-synuclein to specific expression levels. We demonstrate that downregulation of alpha-synuclein in an iPSC-derived neuronal model reduces oxidative stress and shows decrease is mitochondrial DNA damage. Our results provide further proof-of-concept by directly targeting the *SNCA* gene promoter and regulating transcription which present a new experimental model and potentially novel precision medicine for Parkinson’s disease and related alpha-synucleinopathies.

## Results

### S. aureus Cas9 sgRNA design and selection

Previous studies have shown that targeting of a nuclease-dead Cas9 (dCas9) to the promoter region of coding genes lead to robust and specific gene silencing due to interference with the gene transcription machinery. This has been achieved by introduction of inactivating mutations in the Cas9 gene in the two catalytic domains HNH and RuvC, which abolishes the nuclease activity without altering its ability to bind target DNA, blocking the transcriptional process as an RNA-guided DNA-binding complex^20-22^. When dCas9 is fused to one or more KRAB domains, the downregulation is even more pronounced presumably due to epigenetic changes in histone modification with loss of histone H3-acetylation and an increase in H3 lysine 9 trimethylation in the promoter target region^23,24^.

For this study, we chose the *S. aureus* dCas9 (SadCas9) which has a more complex PAM sequence (5’-NNGRRT-3’) compared to *S. pyogenes* Cas9 (5’-NGG-3’) but is smaller in size (1053aa versus 1368aa) and more suitable for downstream *in vivo* applications e.g. when applied by gene therapy vector. We hypothesized that guiding SadCas9 to the promoter region of *SNCA* could downregulate alpha-synuclein in human iPSC-derived neuronal cultures. We followed sgRNA design criteria of targeting dCas9-KRAB to a window of DNA from −50 to +300 bp relative to the transcriptional start site (TSS) in the promoter, assuming a maximum in the ∼50–100 bp region downstream of the TSS^25^. Specifically, for sgRNA design in the *SNCA* promoter we selected a region comprising approximately 3kb upstream of the translation start site in exon 2 (GRCh37/hg19; chr4:90,756,651-90,759,650) including the three TSS for the *SNCA* gene. The CRISPOR online tool^26^ predicted 101 sgRNAs in the *SNCA* promoter containing the 5’-NNGRRT-3’ PAM sequence for SadCas9 irrespective of regulatory control regions. We then selected for our *in vitro* screen in HEK293T cells 32 sgRNAs based on the following selection criteria: 1) absence of termination signals, 2) proximity to TSS -50bp/+300bp, and 3) when two sgRNAs were superimposed (e.g. forward and reverse strand), the sgRNA with higher CRISPOR score was selected (***Figure 1, Supplementary Table 1***). Off-target analysis was performed by CRISPOR tool **(*Supplementary Material, Table 1*)** and is summarized in ***Table 1*** for sgRNAs that we selected for further functional testing in human iPSC cultures.

**Figure 1:**
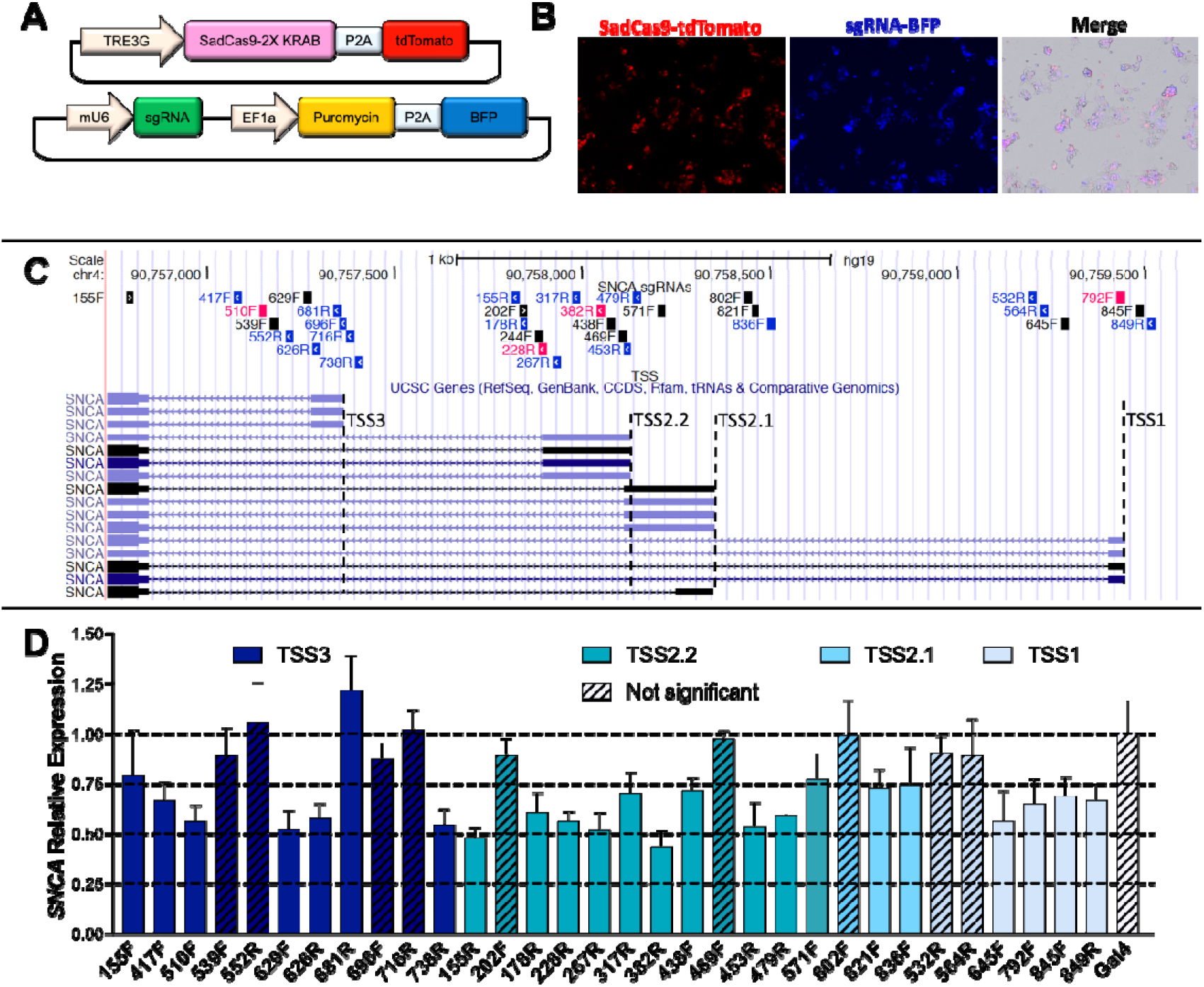
Screening of sgRNAs targeting different TSSs in the *SNCA* promoter region. (**A**) Schematic representation of expression vectors for inducible-SadCas9 and sgRNAs. (**B**) Transient transfection in HEK293T. Cells were co-transfected with 3 plasmids (pHR:pTRE3G-SadCas9-2xKRAB-p2a-tdTomato, pCMV-rtTA, and pHR:mU6-sgRNA-EF1A-Puro-p2a-BFP) and visualized 24 h post-doxycycline treatment. (**C**) Genome browser view of *SNCA* promoter region showing sgRNAs targeted to different TSS within ∼3kb upstream from translation start site in exon 2. sgRNAs in black target + strand (F), while sgRNAs in blue target – strand (R). sgRNAs in pink were selected for experiments in hiPSCs. (**C**) *SNCA* mRNA expression in HEK293T cells transiently transfected with SadCas9 and sgRNAs. Relative expression of *SNCA* mRNA was measured by qPCR and normalized by expression of GAPDH gene. Calibrator sample is sgRNA against the prokaryotic gene *gal4* (white). Data are displayed as mean ± SD for 3 independent transfections (n = 3). Differences between groups were detected by ANOVA with Dunnet post-test. Etched bars indicate non-significant difference in comparison to control *gal4*, TRE, tetracycline responsive element, BFP, blue fluorescent protein, KRAB, Krüppel-associated box, TSS, transcription start site

### CRISPR interference screen by transient transfection of SadCas9/sgRNA in HEK293T cells

To test the ability of the 32 selected sgRNAs to downregulate *SNCA* mRNA, HEK293T cells were transiently co-transfected with three plasmids: i) tetracycline-inducible SadCas9 tagged with tdTomato (pHR:TRE3G-SadCas9-2xKRAB-p2a-tdTomato), ii) sgRNAs tagged with blue fluorescent protein (pHR:mU6-sgRNA-EF1A-Puro-p2a-BFP), and iii) a trans-activator molecule (pCMV-rtTA), henceforth SadCas9-tdTomato, sgRNA-BFP and rTTA. Transient transfection in HEK293T cells had efficiency of about 65%. Cells were treated with doxycycline for 48h to induce expression of SadCas9. *SNCA* mRNA levels were measured 72h post-transfection using Taqman qPCR. A non-targeting sgRNA (*gal4*) was used as negative control.

We identified 23 sgRNAs that significantly reduced mRNA expression of *SNCA* **(*Figure 1D*)**. Around TSS3 which is closest to the translational start site of the *SNCA* (555 bp upstream of translational start site), four sgRNAs showed 50-60%, and two sgRNAs showed 70-80% of *SNCA* expression compared to non-targeting control. Interestingly, one sgRNA showed an increase *SNCA* mRNA expression by approximately 20% compared to sgRNA *gal4* (dark blue columns, Figure 1D). Around TSS2 (TSS2.1 and TSS2.2 are overlapping) 12 sgRNAs exhibited a significant downregulation, with seven showing 45-55% expression and the remaining five sgRNAs reaching 65-75% compared to the *gal4* sgRNA expression (turquoise and light blue, figure 1D). Lastly, around the most upstream located TSS1, we identified four sgRNAs that downregulated *SNCA* mRNA expression to 60-70% of *gal4* sgRNA control levels (very light blue, Figure 1D). For further characterize the CRISPRi system in human iPSCs and neuronal cultures, we selected sgRNAs with different levels of downregulation in HEK293T cultures (approximately 45% for 382R, 60% for 228R and 510F, and approximately 70% for 792F), and proximity to all TSSs to test in human iPSC.

### CRISPR interference downregulates the expression of alpha-synuclein in patient-derived SNCA-triplication and healthy control iPSCs

Next, we prepared lentivirus for SadCas9-tdTomato and rtTA constructs to generate stable inducible SadCas9 human iPSC lines. We transduced human iPSCs from a patient with an *SNCA* genomic triplication and a mutation-negative healthy sibling^9,27^ with both vectors (***Figure 2A***), henceforth called SadCas9-iPSC. After SadCa9 activation with doxycycline in the iPSC culture, we used fluorescence-activated cell sorting to select only cells that expressed tdTomato indicating lentivirus integration (***Supplementary Figure 1***). Within this pool of tdTomato positive cells, there were heterogenous levels of SadCas9-tdTomato expression probably due to differences in location and number of lentiviral integrations. Since different SadCa9 expression levels might affect the regulation of the target gene, in a subsequent step, we derived clonal iPSC lines by serial dilution and selected clonal iPSC lines with an intermediate expression level of SadCas9 (***Figure 2A***). Clonal iPSC lines remained pluripotent after transduction and selection and expressed pluripotency marker OCT4 (***Supplementary Figure 2***).

**Figure 2:**
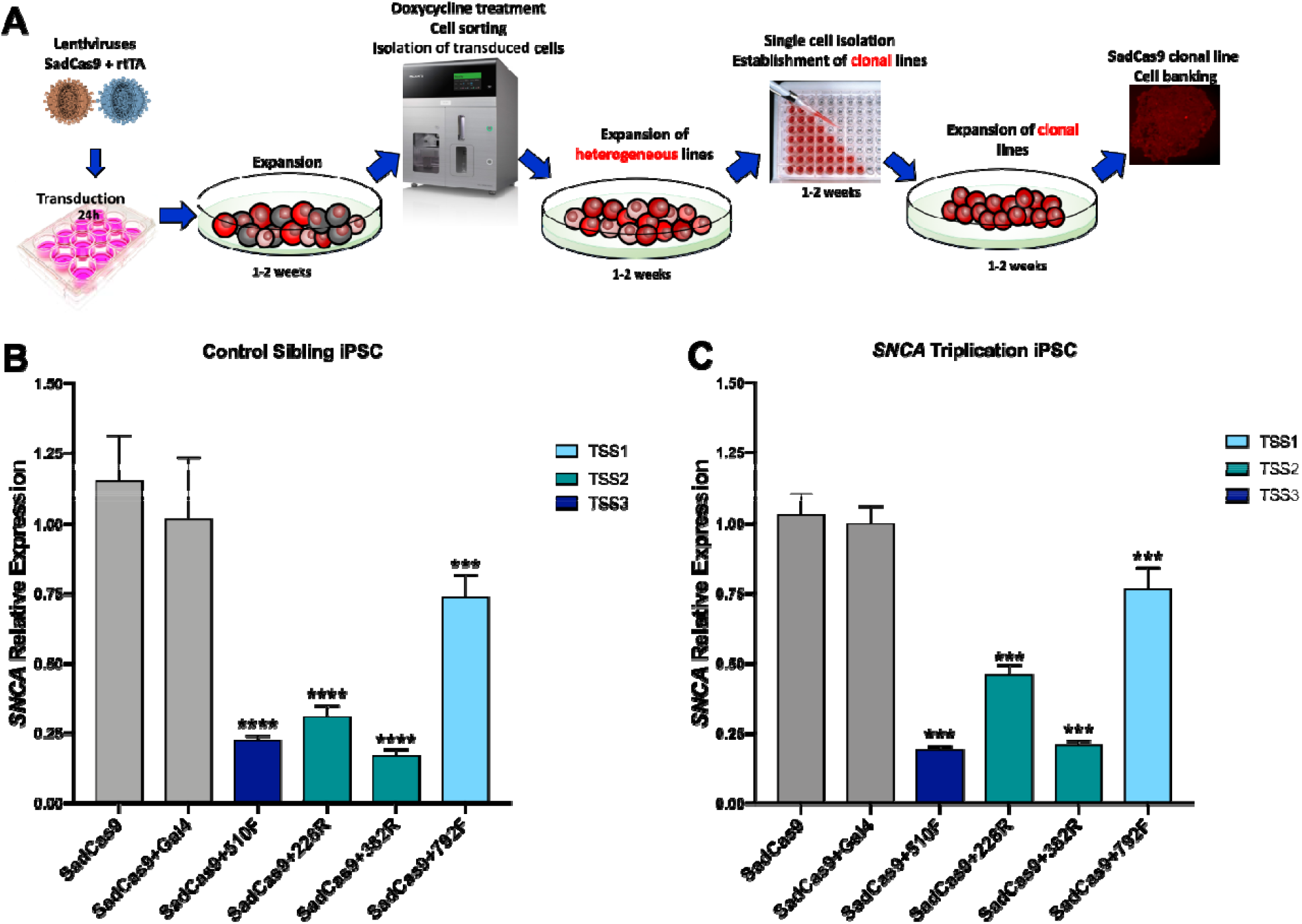
CRISPR/SadCas9-mediated downregulation of alpha-synuclein in patient-derived *SNCA* triplication and healthy sibling control iPSCs. (**A**) Workflow for generation of clonal iPSC lines expressing SadCas9. (**B**) Inhibition of *SNCA* mRNA expression by CRISPRi in *SNCA*-triplication and control sibling (**C**) iPSCs. Clonal SadCas9 iPSC lines were transduced with sgRNAs and isolated by FACS for BFP expression. SadCas9 expression was induced by treatment with 1 μg/mL doxycycline for 48h. Data is represented as mean ± SD for 3 experiments in duplicates (n=6). Differences between groups were detected by ANOVA (p<0.001***, p<0.0001****)

After generating clonal SadCas9-iPSC lines, we tested different sgRNAs identified from the HEK293T cell screen. First, we prepared lentivirus for selected sgRNAs (382R, 228R, 510F, and 792F) using the sgRNA-BFP construct. We then transduced the clonal SadCa9-iPSCs with the sgRNA lentivirus and sorted the cell population for blue fluorescence. It was critical to perform these steps sequentially without introduction of doxycycline during the selection process, as we observed prolonged downregulation of the *SNCA* gene when cells were exposed to doxycycline in the presence of both SadCa9 and sgRNA expression (data not shown). We then exposed the different sgRNA-transduced clonal iPSC cultures to doxycycline for 48h and measured *SNCA* mRNA expression. The level of downregulation was more pronounced when compared to the HEK293T cultures as 100% of cells expressed both the doxycycline-inducible SadCas9 and sgRNA-BFP. In clones from both the patient with the *SNCA* genomic triplication (H4C2) and sibling control (H5C2), two of the SadCas9/sgRNAs showed approximately 25% expression compared to SadCas9/*gal4* sgRNA control (75% downregulation for 382R and 510F), one SadCas9/sgRNA 228R had 50% expression compared to SadCas9/*gal4* sgRNA control (50% downregulation), and one SadCas9/sgRNA 792F had 75% expression level compared to SadCas9/*gal4* sgRNA control (25% downregulation) (***Figure 2 B, C***).

### SNCA isoform downregulation in patient iPSC-derived floorplate progenitors (FPp1)

Besides the full-length mRNA transcript of the *SNCA* gene (*SNCA*140), there are three alternative transcript isoforms that are found in the brain. The canonical *SNCA* isoform contains all 5 coding exons and codes for a protein of 140 amino acid residues (*SNCA*140). *SNCA*126 isoform lacks exon 3, *SNCA*112 isoform is lacking exon 5, and *SNCA*98 isoform is missing both exons 3 and 5. The four *SNCA* isoforms are differentially expressed in the brains of patients with PD and other forms of neurodegeneration^28-32^. Characterization of the aggregation of the *SNCA* isoforms by electron microscopy showed fibril bundles for the *SNCA*140, shorter fibrils for *SNCA*126, and interestingly annular structures for *SNCA*98^33^.

Because alternative promoter usage has been shown to influence mRNA splicing^34,35^, we wanted to determine to what extent the different *SNCA* isoforms would be affected by the CRISPR/SadCas9-mediated downregulation when targeting different transcriptional start sites of the *SNCA* promoter. After optimization of isoform-specific primers (***Supplementary Table 3, Supplementary Figure 3***), we performed mRNA expression analysis for all four isoforms. The level of isoform expression in the floorplate progenitors is lower to the expression in human brain^33^, but the ratios of the shorter isoforms are comparable with about 0.003% expression of the *SNCA*112 compared to *SNCA*140 (set to 100%) and less than 0.001% for the *SNCA*126 and *SNCA*98 isoforms.

The level of *SNCA* downregulation for the four different sgRNAs was directly comparable to the data in clonal iPSC lines for the full-length *SNCA*140 isoform (***Figure 2B, Figure 3***). However, when we measured the smaller isoforms *SNCA*126, *SNCA*112, and *SNCA*98, only the two sgRNAs (382R and 510F) mediated significant downregulation, whereas expression of the other two sgRNAs (228R and 792F) were not significantly different for the smaller *SNCA* isoforms (***Figure 3***).

**Figure 3.**
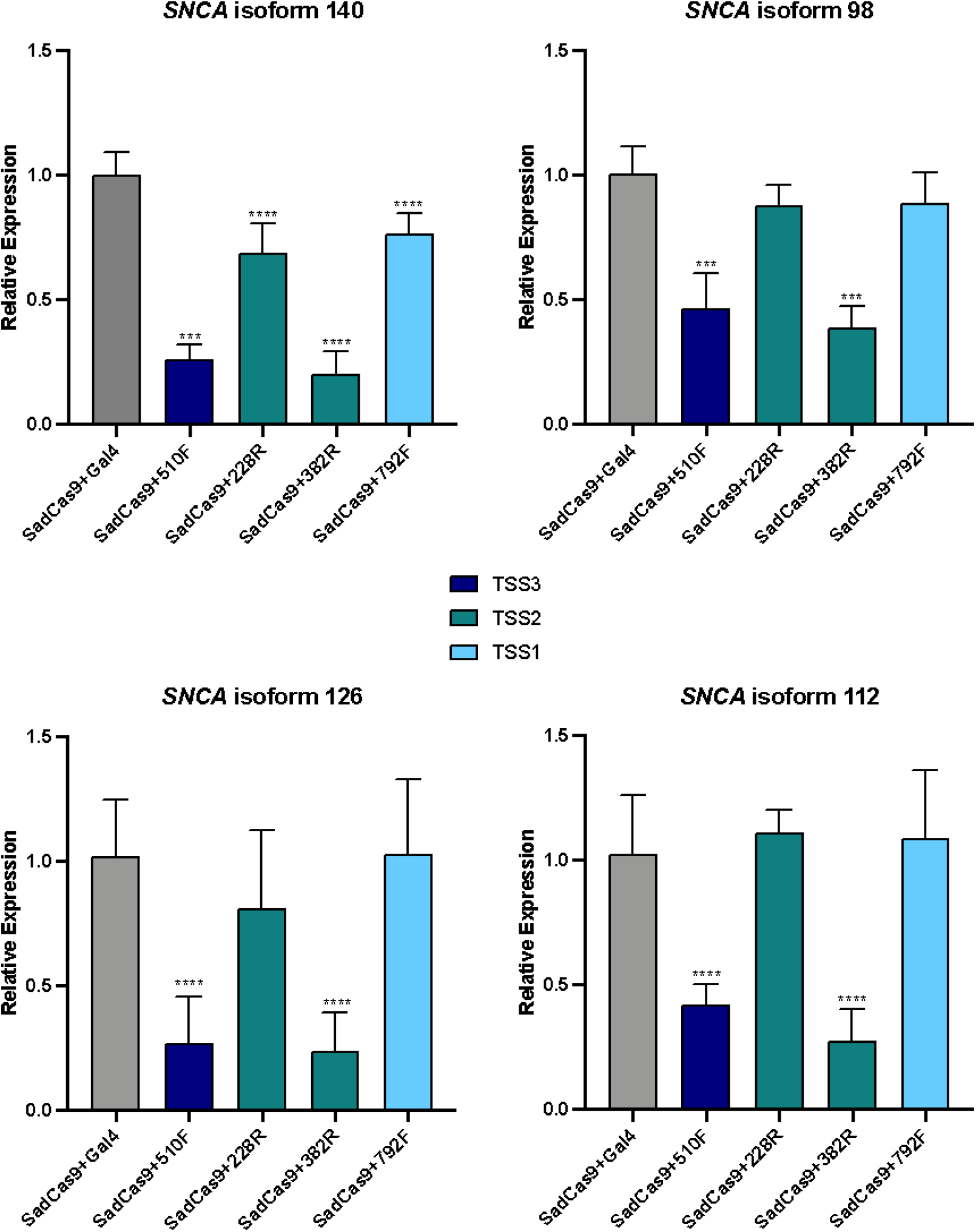
*SNCA* isoform downregulation in iPSC-derived floorplate progenitors (FPp1) from patient with *SNCA* triplication. Isoform expression levels in clonal *SNCA* triplication FPp1s post treatment with doxycycline 1 µg/mL for 5 days prior to cell harvest. Relative *SNCA* isoform expression of FPp1s is measured by qPCR and normalized to GAPDH expression. Calibrator sample is sgRNA against the prokaryotic gene *gal4*. Data are displayed as mean ± SD for 2 independent biological experiments with 3 technical replicates (n=6). Differences between groups were detected by ANOVA (p<0.0005***, p<0.0001****).

### CRISPRi-mediated alpha-synuclein downregulation reduces ROS and lipid peroxidation in neuronal cultures from SNCA triplication carrier

A common PD phenotype is increase in reactive oxygen species (ROS) damage^36-38^ which can also cause damage to mitochondrial (mt)DNA because of the proximity of mtDNA to ROS production at the inner mitochondrial membrane and the lack of protection afforded by histones^39^. We performed different assays related to oxidative stress which show that alpha-synuclein downregulation reduces the effects of oxidative stress (***Figures 4***). We differentiated the four clonal *SNCA* CRISPRi iPSC lines into floorplate progenitors (***Supplementary Figure 4)***. On day 15 of the differentiation process, we exposed cells to doxycycline at 1ug/ml to activate SadCas9 expression.

**Figure 4.**
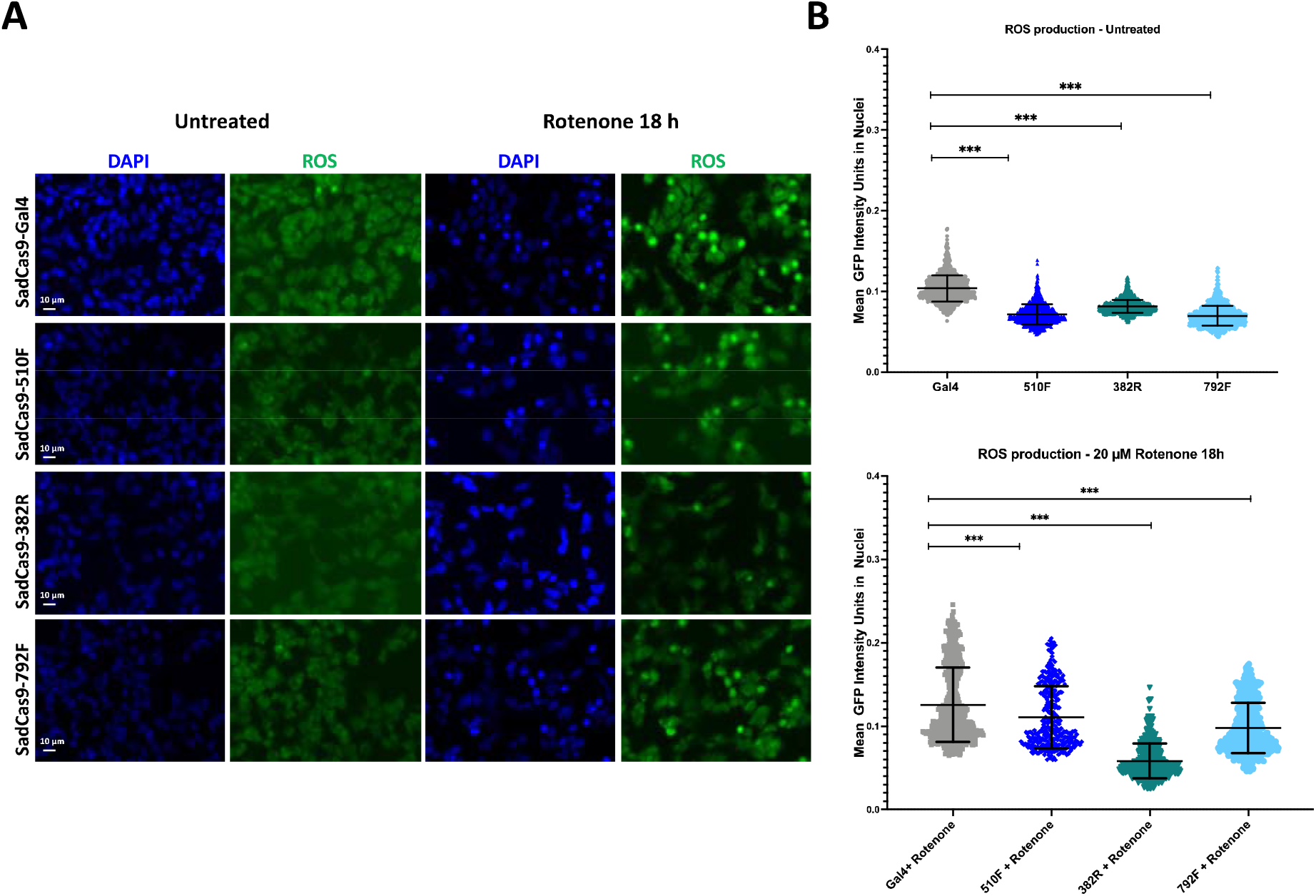
CellROX® Green Oxidative Stress assay in iPSC-derived floorplate progenitors from patient with *SNCA* triplication. The figure illustrates measurement of reactive oxygen species (ROS) using a fluorogenic probe that presents with a strong fluorogenic signal upon oxidation and localizes to nuclei. (**A**) Representative images of reactive oxygen species (ROS), (**B**) Mean intensity of ROS per nuclei in rotenone treated cells. The upper panel represent the untreated naïve condition, and lower panels represent 18 h rotenone treatment (n=9 images per condition; 5,957 nuclei analyzed). Differences between groups were detected by ANOVA (p<0.001 ***).

The ROS assay is based on the CellROX® green oxidative stress reagent, which is a fluorogenic probe that measures reactive oxygen species (ROS) in live cell cultures. The green cell-permeable DNA dye is weakly fluorescent while in a reduced state and upon oxidation exhibits a strong nuclear fluorogenic signal.

Five days after doxycycline treatment to induce SadCas9 expression, we performed the CellROX® Green Oxidative Stress assay in iPSC-derived neuronal cultures carrying the *SNCA* triplication. We detected a decrease in steady-state ROS production in cells with SNCA expression downregulated via CRISPRi in comparison to non-targeting control. We then treated these cultures with rotenone to induce mitochondrial damage and observed all sgRNAs tested significantly reduced the ROS production compared to control. The effect seems to be more pronounced for the sgRNA that shows the stronger downregulation (382R, 75% *SNCA* mRNA knockdown).

### Rescue of mitochondrial (mt)DNA damage after CRISPR-guided a-syn downregulation iPSC-derived neuronal cultures from patient with SNCA triplication

Oxidative damage may lead to mtDNA damage, mutations and degradation, or directly trigger cell death^40^ and plays a role in aging and neurodegeneration^41^. The mitochondrial genome is particularly susceptible to accumulation of oxidative damage, which is a particular problem for the brain because neurons are post-mitotic and long-lived.

We applied the Mito DNA_DX_ assay, a robust quantitative real-time assay of mtDNA damage in a 96-well platform^42^. In brief, less PCR product will be produced when mtDNA damage or lesions block the ability of the DNA polymerase to replicate and the PCR amplification of a large fragment specific to the mitochondrial genome is analyzed^43,44^ Thus, mtDNA damage or repair slowing down or impairing DNA polymerase progression will be detected.

Five days after Cas9 expression was induced in iPSC-derived floorplate progenitors, we collected cell pellets and performed the Mito DNA_DX_ assay. SgRNA 382R that downregulates *SNCA* expression by 75% showed significant reduction in mtDNA lesions compared to the control sample. The other three sgRNAs that show less downregulation do not promote decrease in mtDNA damage (***Figure 5A***). However, mtDNA copy number did not vary between samples (***Figure 5B***).

**Figure 5.**
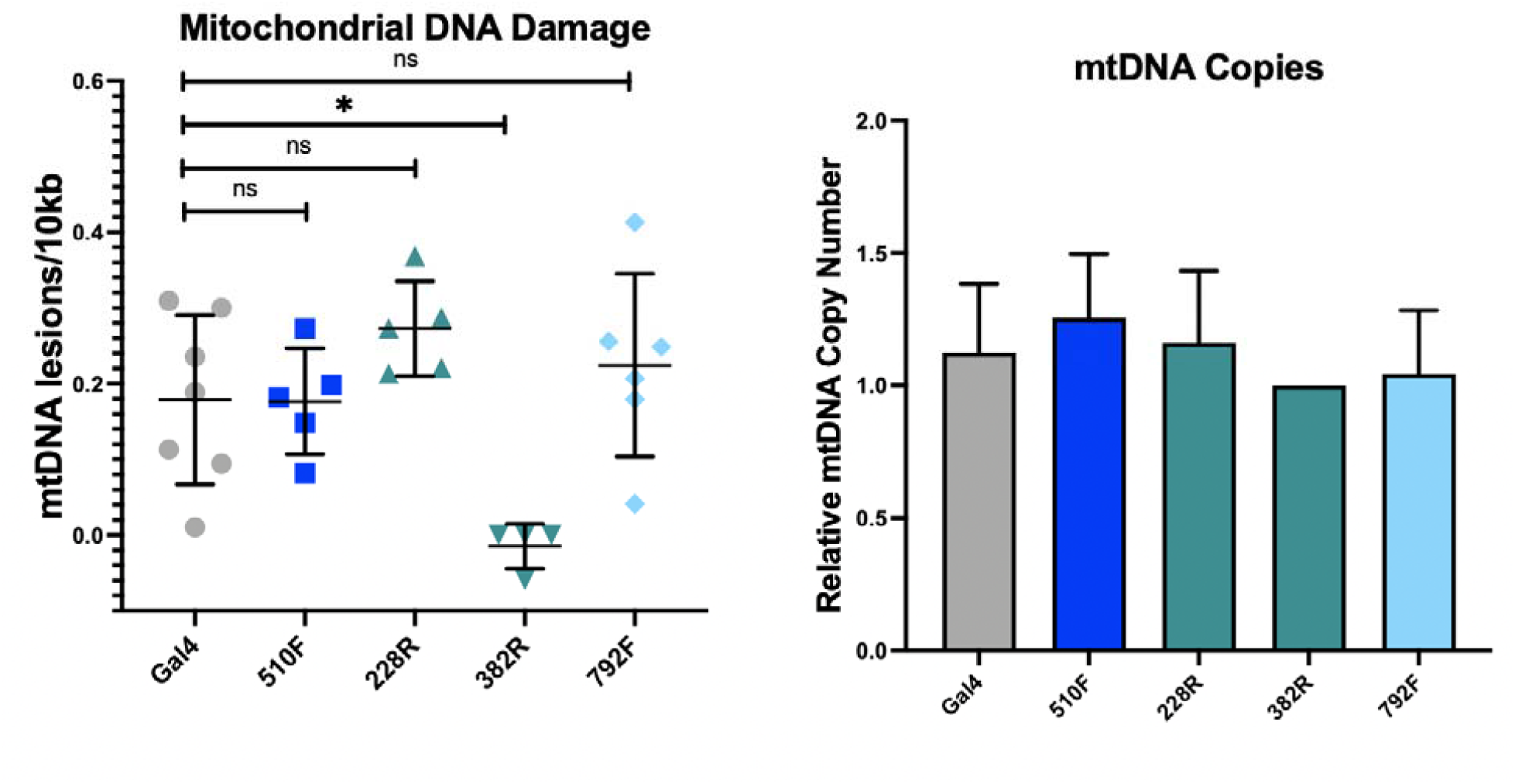
Mitochondrial DNA damage ameliorated in neuronal cultures from *SNCA* triplication carrier with CRISPR-guided *SNCA* downregulation. A. sgRNA382R shows significant reduction in mtDNA lesions compared to *gal4* control (p<0.05). All other sgRNA did not show significant improvement of mtDNA damage (n=4 to 7 biological replicates). B. mtDNA copy numbers did not vary between samples.

## Discussion

Alpha-synuclein has become a leading target for development of new therapeutic strategies aimed at disease modification for PD and related synucleinopathies. Interest in alpha-synuclein as a therapeutic target has been greatly enhanced by the knowledge that increased expression of wildtype alpha-synuclein protein alone can lead to neurodegeneration, as observed in patients with duplications and triplications of the *SNCA* genomic locus and also common genetic promoter and non-coding variants of the *SNCA* gene (e.g. Rep-1 allele) which may upregulate alpha-synuclein expression ^7,9,45-49^. It is thought that the mere overexpression of alpha-synuclein over time can lead to neurodegeneration and neuronal demise exemplified by these genetic studies. Especially the gene dosage effect of alpha-synuclein expression and severity of symptoms, age at onset and disease progression provides confidence about causality.

These clinical genetic observations and pre-clinical *in vitro* and *in vivo* studies have led to the premise that lowering the alpha-synuclein content and/or eliminating toxic alpha-synuclein species in cells could be the key to slowing, reversing, or even preventing the disease. Many different strategies have been proposed from reducing alpha-synuclein mRNA by RNA interference or antisense oligonucleotides, inhibiting alpha-synuclein aggregation, promoting intracellular degradation of alpha-synuclein, increase of extracellular alpha-synuclein degradation through active or passive immunotherapies to reduction of uptake of extracellular alpha-synuclein^4,50^. Unfortunately, recent clinical trials using alpha-synuclein antibodies failed their primary clinical endpoints and such programs were discontinued^51^.

While reduction of extracellular alpha-synuclein might not be sufficient to influence clinical disease course of PD or related alpha-synucleinopathies, it is critical to develop other strategies to modulate alpha-synuclein expression to understand the effects of different gene dosage levels and to expand the armamentarium of alpha-synuclein regulation for future therapeutic use.

CRISPR-mediated gene engineering has become an attractive tool and potential therapeutic platform for gene therapies. For alpha-synuclein, using CRISPR knock-out strategies, several *SNCA* knockout lines have been developed^52,53^. Another approach for *SNCA* downregulation uses targeted editing of DNA methylation of the *SNCA* gene promoter region which shows a downregulation of alpha-synuclein by 30%^54^.

Our study is built on previous data that showed evidence for proof of concept that the *SNCA* promoter can be successfully targeted by CRISPR/Cas9 interference^19,52^. The group found one sgRNA in the region of the transcriptional start site 2 (TSS2) of the *SNCA* promoter that showed a 50% *SNCA* downregulation in HEK293 cultures and human iPSC-derived neurons. In that study, the *S. pyogenes* version of CRISPR/Cas9 was used which is 4.2 kb in length. Here, we used *S. aureus* dCas9 guided to the promoter region of *SNCA* gene which also robustly mediated reduction of mRNA and protein levels *in vitro. S. aureus* Cas9 is one of the smaller Cas9 proteins which are developed for *in vivo* gene therapy application. Even smaller Cas9 proteins have been discovered and characterized such as CjCas9^55^ or CasX^56^, but their PAM recognition sequences can be complex, thus lowering the yield of predicted sgRNAs.

Although the PAM sequence for the SaCas9 is more complex, we identified over 100 sgRNAs in the 3kb promoter region of the *SNCA* gene which we narrowed down to 32 for the HEK293T screen and 22 significantly reduced *SNCA* mRNA expression.

Several smaller splice isoforms of the *SNCA* gene *have* been described and characterized^57^. There is some evidence that *SNCA*112 and *SNCA*98 promote alpha-synuclein seeding whereas *SNCA*126 seems to be protective^29,58^. Because of these unique and possibly disease-relevant features, we also tested to what extent the expression of *SNCA* isoforms have been affected by the SadCas9-mediated knockdown. Since alternative promoter start sites can have an effect on mRNA splicing, e.g. for the MAPT gene^34^ - where alternative promoter usage leads to novel shorter transcripts in Alzheimer disease - we wanted to test if any of our selected sgRNAs targeting different transcriptional start sites would affect the downregulation or expression ratio of the different isoforms. All sgRNAs downregulated *SNCA* expression for the most abundant full-length *SNCA*140 form. Two of the sgRNAs downregulated all four tested *SNCA* isoforms, whereas the two sgRNAs that mediated less downregulation, did not affect expression levels of any of the smaller isoforms. This data demonstrates CRISPRi ability to selectively modulate different mRNA isoforms, opening the exciting possibility for future work to identify sgRNAs selectively sparing *SNCA*126 expression which is thought to be protective in the context of PD^32^. Characterization of SNCA protein levels in response to varying degrees of CRISPRi downregulation would further elucidate the relationship between SNCA isoforms and their translational potential.

Even though a lot of effort has been put into lowering the amount of alpha-synuclein by various technologies, there is still the open question of what is a safe degree of alpha-synuclein knockdown. We think that pre-clinical models of human induced pluripotent stem cell-derived neurons could help answering this question. This model has several advantages: first, it is based on a human background and the model has been derived from a patient with an gene copy number mutation of the *SNCA* gene^27^, expresses twice as much alpha-synuclein protein, shows robust cellular phenotypes for increased oxidative stress and impaired energy metabolism^59,60^, and clinically showed a severe form of Lewy body disease^9,61^. Second, inducible SadCas9 expression in mature neurons can mimic a therapeutic intervention in an adult organism, whereas CRISPR knockout iPSC lines for the *SNCA* genomic locus^53^ can elucidate the developmental importance of alpha-synuclein^62,63^.

Previously, we reported that lymphoblastoid cell lines and iPSC-derived neuronal cultures from *LRRK2* p.G2019S mutation carriers have increased mtDNA damage^64,65^. When we genetically corrected the *LRRK2* mutation or treated the cultures with a LRRK2 kinase inhibitor mtDNA damage was no longer detectable. Thus, we concluded that the mtDNA damage phenotype can be unambiguously attributed to the *LRRK2* p.G2019S mutation.

We also hypothesized a similar mtDNA damage phenotype also occurs for the *SNCA* genomic triplication as we have shown that neuronal cultures from an *SNCA* triplication case shows higher levels of oxidative stress ^27,59^ and should also lead to mtDNA damage. We hypothesized that lowering a-syn expression by CRISPR-guided downregulation might improve mtDNA damage. However, only with a 75% downregulation of *SNCA* mRNA expression in the *SNCA* triplication, we achieved a significant rescue of mtDNA damage. This indicates that alpha-synuclein intracellular levels need substantial alteration to show a therapeutic effect.

In summary, CRISPR/Cas9 gene-engineering technologies with the power to precisely target genomic locations have the potential to translate into precision medicine^66^. Our studies show proof-of-concept in vitro that *SNCA* mRNA reduction leads to a functional rescue of pathophysiological phenotypes related to PD and neurodegeneration. Further studies *in vivo* will be critical to develop a translational program and show proof whether this approach could be a viable therapeutic strategy.

## Methods

### Cloning of CRISPR/sgRNA lentiviral constructs with fluorescent selection markers

A tetracycline-inducible promoter (TRE3G) was used to control the expression of *S. aureus* dCas9 in a lentiviral vector. To facilitate selection of cells by FACS, pHR:TRE3G-SadCas9-2xKRAB-p2a-tdTomato was subcloned from a pHR:TRE3G-SadCas9-2xKRAB-p2a-zeo (A gift from Professor Stanley Qi), where zeocin resistance gene (zeo) was replaced by a fluorescent marker, tdTomato. Briefly, backbone was digested with BamHI and NotI to linearize vector.

SadCas9-2xKRAB-p2a segment was amplified by PCR from backbone using high-fidelity polymerase (Phusion, NEB, Cat. No. M0530S); and tdTomato was PCR amplified from a proprietary expression vector using high-fidelity polymerase. Assembly PCR primers were designed to include at least 40 bp of homology with adjacent fragments. Digested backbone and PCR amplified fragments were joined via seamless cloning (NEBuilder® HiFi DNA Assembly Master Mix, NEB, Cat. No. E2621S). Assembled plasmid was Sanger-sequenced to ensure correct assembly and reading frame. The reverse tetracycline-controlled transactivator was expressed from pCMV-rtTA (rtTA-N144), a gift from Andrew Yoo (Addgene # 66810).

sgRNAs were restriction enzyme-cloned into an expression vector tagged with blue fluorescent protein, pHR:mU6-sgRNA-EF1A-Puro-p2a-BFP, a gift from Professor Stanley Qi. Briefly, backbone was linearized using BstXI and XhoI, and subsequently dephosphorylated with 1U of Calf Intestinal Alkaline Phosphatase (ThermoFisher, Cat. No. 18009019), following manufacturer’s protocol. sgRNAs were generated using PCR amplification with primers containing the appropriate restriction sites **(Supplementary Table 2)**. sgRNA PCR products were digested with BstXI and XhoI to create compatible ends. Linearized vector and sgRNAs were ligated using T4 ligase (ThermoFisher, Cat. No. EL0014).

### sgRNA design with CRISPOR tool

We used CRISPOR tool^26^ for the sgRNA design and selection. We divided the *SNCA* promoter region into three 1kb segments spanning 3kb upstream exon 2 (GRCh37/hg19; chr4:90,756,651-90,759,650) due to CRISPOR sequence size limitations. The *SNCA* promoter comprises three transcriptional start sites (TSS) and sgRNAs were designed at positions - 50 to + 300 bp around each of the three TSSs. Designed *S. aureus* Cas9 sgRNAs are 21-bp long and are adjacent to a PAM spacer sequence 5’-NGGRRT-3’ **(*Supplementary Table 1*)**. As a negative control, we used a sgRNA against a gene from *Saccharomyces cerevisiae, gal4*, which does not target the human genome.

### Cell culture and neuronal differentiation

#### HEK293T cell culture

Cells were maintained in DMEM:F12 medium (ThermoFisher, Cat. No. 11320033) supplemented with 10% tetracycline-free FBS (Takara Bio, Cat. No. 631367), 100 U/mL Penicillin-Streptomycin (ThermoFisher, Cat. No. 15140122) and 1X MEM Non-Essential Amino Acids Solution (ThermoFisher, Cat. No. 11140050). Medium was changed every two days and cells were passaged by enzymatic treatment with TrypLE (ThermoFisher, Cat. No. 12604021) when 90-100% confluent. Cells were subcultured at 1:6 ratio.

#### Human iPSC culture

Human iPSCs were from a patient with *SNCA*-triplication (Iowa Kindred) and from a control sibling^9,27^. iPSCs were maintained under feeder-free conditions as colonies on 12-well plates coated with Geltrex™ (ThermoFisher, Cat. No. A1413302) diluted 1:70, using StemFlex™ medium (Thermo Fisher, Cat. No. A3349401) supplemented with 100 U/mL Penicillin-Streptomycin (ThermoFisher, Cat. No. 15140122). Confluent iPSC colony plates were passaged manually once a week. Manual passaging consisted of slicing the colonies into small square pieces with a 25-gauge needle and then pipetting the colony pieces onto a new plate Geltrex-coated plate. After each passage, cells were maintained for 24 h in medium supplemented with 1µM RHO/ROCK pathway inhibitor Thiazovivin, THZ (ReproCell, Cat. No. 04-0017).

For experiments requiring exact cell numbers, iPSC colonies were dissociated using enzymatic treatment with Accutase (Innovative Cell Tech., Cat. No. AT104-500). Accutase was applied for 5 min until cells start to detach and cells were seeded at a subcultivated at a ratio of 1:6. Dissociated iPSCs were maintained as monolayers on plates coated with 10 µg/µL rhLaminin-521 (ThermoFisher, Cat. No. A29248). After each passage, cells were maintained for 24 h in medium supplemented with 1µM THZ. iPSCs maintained as monolayers were propagated every 5 to 6 days. For SadCas9 activation, upon reaching confluency, cells were treated with 1µg/mL doxycycline for 48 h and then collected for downstream experiments.

#### Floor plate progenitor differentiation

iPSCs were maintained as monolayers on plates coated with 10 µg/µL rhLaminin-521 in StemFlex medium, as described above. iPSCs were differentiated into midbrain floor plate progenitors via the PSC Dopaminergic Neuron Differentiation Kit (ThermoFisher, Cat. No. A3147701) in a stepwise fashion comprising two phases: specification and expansion. Specification medium induces iPSCs to differentiate towards midbrain lineage floor plate progenitors (FPp). Expansion medium promotes proliferation of FPps.

On day -1 of differentiation (plating day), cells were passaged using Accutase as described above and seeded at a density of 5.56 × 10^4^ cells /cm^2^ in 6-well plates coated with 10 ug/Ml rhLaminin-521 in StemFlex medium supplemented with 1µM THZ. On the next day (Day 0), cells reached 20-40% confluence, and medium was replaced with specification medium, following manufacturer’s instructions. Culture media was changed on Days 3, 5, 7 and 9 of differentiation. On Day 10 of differentiation, FPp cells were enzymatically passaged with Accutase and seeded at 1:2 ratio in 2 mL of expansion medium supplemented with 2 µM THZ onto 6-well plates that were sequentially coated first with Geltrex and then with 15 ug/mL mouse laminin (ThermoFisher, Cat. No. 23017015). Medium was replaced on Day 11 to remove THZ. When cells reach 100% confluency on Day 12-13, FPp cells were passaged and seeded at 1.67×10^5^ cells /cm^2^ in 6-well plates using expansion medium supplemented with 2 µM THZ. Culture media is replaced every other day. For all experiments, doxycycline was added on day 12 for 5 days. Cells were collected and experiments were carried out between day 17-20.

### Transient transfection

HEK293T cells were seeded on 6-well plates at density of 2.22 × 10^4^ cells/cm^2^ overnight. At 80-100% confluency, cells were transfected with 1µg of each plasmid: pHR:TRE3G-SadCas9-2xKRAB-p2a-tdTomato, pCMV-rtTA and pHR:mU6-sgRNA-EF1A-Puro-p2a-BFP (1:1:1), using 25 µL TransIT®-LT1 Transfection Reagent (Mirus Bio, Cat. No. MIR 2300) in a final volume of 1mL DMEM medium supplemented with 10% tetracycline-free FBS. After 24 h, medium was changed to DMEM:F12 medium supplemented with 500 ng/mL doxycycline to activate SadCas9 expression. Cells were collected for RNA extraction 72 h post-transfection/48 h doxycycline treatment. Experiments were performed in triplicates.

### Lentivirus production

For lentiviral production, we used 2^nd^ generation lentiviral packaging plasmid pCMVR8.74 (Addgene #22036) with envelope-expressing plasmid pMD2.G-VSV-G (Addgene #12259). HEK293T cells were plated 24 h prior to transfection in 10 cm plates at density of 2 × 10^4^ cells/cm^2^. Plasmid DNA ratio was <u>9:8:1</u> for <u>transfer</u> (pHR:TRE3G-SadCas9-2xKRAB-p2a-tdTomato or pCMV-rtTA or pHR:mU6-sgRNA-EF1A-Puro-p2a-BFP) : <u>pCMVR8.74</u> : <u>pMD2.G-VSV-G</u> respectively, in a total of 10ug DNA. A final volume of 5 mL with 25 µL TransIT®-LT1 Transfection Reagent was used for transfection. Medium was changed 24h post-transfection.

Supernatant containing lentiviruses was collected 48 h post-transfection, filtered with a 40 µm filter, and ultracentrifugated for 1.5 h at 25,000 rpm/107,000 g (Sorvall™ WX+, ThermoFisher) to concentrate virus. Viruses were resuspended in 75 µL PBS, aliquoted and stored at -80°C.

### Lentiviral transduction of human iPSCs

Human iPSCs were plated 24 - 48 h before transduction at density of 1.25 × 10^5^ cells/cm^2^ in 12-well plates. Cells were infected 12.5 µL of concentrated lentivirus pTRE-SadCas9-2xKRAB-dtTomato and 12.5 µL of concentrated lentivirus pCMV-rtTA in 400 µL of StemFlex medium supplemented with 6 ug/mL of polybrene (Millipore, Cat. No. TR-1003-G). After 24 h, media was replaced with StemFlex medium and changed every other day until cells reached 100% confluency. Confluent cells were expanded into 6-well plates for 1-2 weeks prior to FACS sorting.

Clonal iPSC-SadCas9 were plated 24 - 48 h before transduction at density of 1.25 × 10^5^ cells/cm^2^ in 12-well plates. Cells were infected 25 µL of concentrated lentivirus pHR:mU6-sgRNA-EF1A-Puro-p2a-BFP in 400 µL of StemFlex medium supplemented with 6 ug/mL of polybrene. After 24 h, media was replaced with StemFlex medium and changed every other day until cells reached 100% confluency. Confluent cells were expanded into 6-well plates for 1-2 weeks prior to FACS sorting.

### Fluorescence activated cell sorting (FACS)

Expanded SadCas9 transduced iPSCs were treated with 1µg/mL doxycycline for 24 h prior to FACS. Briefly, on the day of sorting, 2-10 × 10^6^ cells were dissociated using enzymatic treatment with Accutase for 5 min, centrifuged at 1000 rpm for 3 min, and re-suspended in 1-2 mL of cold PBS. Cell suspensions were filtered through a 100 µM nylon cell strainer (ThermoFisher, Cat. No. 07-201-432) before acquisition on a flow cytometer (Sony, Model SH800). Sorter was equipped with three excitation lasers: 488 nm, 405 nm, and 561 nm; sorter channels used were FL 450/50, FL 525/50, and FL3 600/50.

Cells were sorted based on expression of tdTomato as a marker for expression of SadCas9. Using the 100 μm sorting chip at 50 kHz, an average threshold of 10,000 events per second, cells were sorted for FL3/tdTomato and FL2/PE (for auto compensation) with ultra-purity setup. Sorted cells were transferred to 48-well plates and cultured as described above. Sorted cells were a heterogeneous population showing different levels of SadCas9 expression.

No doxycycline treatment was used prior to sorting of sgRNA-iPSCs. On day of sorting, 2-10 × 10^6^ SadCas9-sgRNA transduced iPSCs cells were dissociated using enzymatic treatment with Accutase for 5 min, neutralized with PBS, centrifuged at 1000 rpm for 3 min, and re-suspended in 1-2 mL of cold PBS.

Cells were sorted based on expression of BFP as a marker for expression of sgRNAs. Using the 100 μm sorting chip at 50 kHz, an average threshold of 10,000 events per second, cells were sorted for FL1/BFP and FL2/EGFP (for auto compensation) with ultra-purity setup. Sorted cells were transferred to 24-well plates and cultured as described above.

### Single cell cloning of SadCas9-iPSCs

To achieve homogeneous levels of SadCas9 expression, SadCas9-positive iPSCs with various expression levels of SadCas9 were clonally selected using serial dilution. A final dilution of 0.5 cells/100 µL of medium was plated into 96-well plates (100 µL/well). Media was changed every 2-3 days for approximately 2 weeks until colonies originated from single cells (clonal) which were further expanded and cryopreserved.

#### Immunofluorescence

For staining of iPSC-derived FPps, rabbit anti-LMX1A (Abcam, Cat. No. ab31006) and mouse anti-FOXA2/ HNF-3 beta (Santa Cruz Bio., Cat. No. sc-101060) were used at 1:500 and 1:200 dilutions, respectively. Secondary antibodies were purchased from ThermoFisher: Goat Anti-Mouse Alexa Fluor 488 (Cat. No. A-10680) and Goat Anti-Rabbit Alexa Fluor 594 (Cat. No. A-11037), both at 1:500 dilution.

Plates were washed with DPBS (ThermoFisher, Cat. No. 14040133) to remove media and debris. Cells were then fixed with 4% paraformaldehyde/DPBS solution for 10 min, followed by three washes with DPBS. Cells were permeabilized with 0.1% Triton-X100 solution for 5-10 min, followed by one wash with PBS. Cells were blocked with blocking buffer containing 5% goat serum (ThermoFisher Cat. No. 16210064) in DPBS for 1 h, followed by three washes with PBS. Primary antibody was added in appropriate concentration and incubated overnight at 4°C. On the next day, all wells were washed three times with DPBS and secondary antibody was added and incubated on a rocker for 1 h at room temperature. Cells were washed with DPBS three times and a Hoechst 33342 (ThermoFisher Cat. No. H3570, 1ug/mL) nuclear stain solution was added to each well and incubated for 5-10 min. Cells were washed three times with PBS and stored at 4°C before fluorescent imaging.

### RNA extraction, cDNA synthesis, and qPCR

Cell pellets were collected by enzymatic dissociation with Accutase, washed once in 2 ml PBS, and stored at -80°C freezer until use. RNA extraction and DNase treatment were performed using the PureLink™ RNA Mini Kit (ThermoFisher, Cat. No. 12183025) following manufacturer’s guidelines. RNA concentration and quality were determined using the NanoDrop Technologies ND-1000 spectrophotometer (ThermoFisher). cDNA synthesis was completed using 1 ug of RNA as input for random primer reaction (High Capacity cDNA Reverse Transcription Kit; ThermoFisher, Cat. No. 4368814) following manufacturer’s guidelines. cDNA was diluted with nuclease-free water to a concentration of 10 ng/µL. For qPCR, 10 ng of cDNA was used as template. TaqMan® Gene Expression Master Mix (ThermoFisher, Cat. No. 4369016) and TaqMan probes for FAM-*SNCA* (ThermoFisher, Assay ID: Hs00240906_m1) and VIC-*GAPDH* (ThermoFisher, Assay ID: Hs99999905_m1) were used to amplify cDNA. qPCR reactions were run in CFX96 Real Time System thermal cycler (Bio Rad) and Cq values were obtained using built-in software. No template control (NTC) samples were included in each plate. RT-control samples were included for each round of RNA extraction and consisted of a cDNA reaction without addition of reverse transcriptase. NTC and RT controls did amplify in any of plates included in the data presented. Relative mRNA expression was calculated using 2^-∆∆Ct^ method ^67^ using *GAPDH* as a reference gene and Excel software for calculations. The calibrator sample was sgRNA-gal4, a sgRNA that does not target any human gene. All samples were run in three technical qPCR replicates. A minimum of two biological replicates were used for each sgRNA tested. Cq outliers were eliminated if SD >0.5 Ct.

### Functional Cellular Assays in iPSC-derived midbrain progenitors

For all fluorescence-based functional assays, iPSC-derived midbrain floor-plate progenitors (FPp) cells were treated with 1 µg/mL doxycycline on day 12 of differentiation. Cells were passaged and seeded at density of 1.43 × 10^5^ cells/cm^2^ into 8-well chamber glass slides on day 14 of differentiation. On day 15 of differentiation, cells were treated with 20 μM rotenone for 18 h. After treatment, cells were stained with functional assay stains. Manufacturer’s instructions were followed for all assays, unless otherwise stated. After staining, cells were fixed in 10% buffered neutral formalin for 15 min and washed with PBS. Nuclei were stained with 1 µg/mL Hoechst 33342 for 1 min. Cells were mounted with glass coverslips using ProLong™ Gold Antifade Mountant (ThermoFisher, Cat. No. P36930) and imaged using a fluorescence microscope (BZ-X700, Keyence). Image processing was performed using built-in image processor.

#### CellROX® Green Oxidative Stress staining

(Thermo Fisher, Cat. No. C10444): CellROX® Green Reagent is a DNA dye, thus strong nuclear signal indicates higher production of reactive oxygen species. We treated the cells with 20 µM rotenone for 18 hrs. Culture medium was removed and replaced with HBSS solution containing 5 μM of CellROX® Reagent and incubated for 30 min at 37°C. Cells were incubated at 37°C for 30 minutes, then fixed with 10% buffered neutral formalin for 15min. Hoechst was used as nuclear counterstain. Excitation and emission were 485/520nm (Green). Fluorescence was measured within 24 h after staining.

#### Quantifying mtDNA damage with a PCR-based assay

DNA isolation and quantification was performed as previously described^43,44,68^ using a high molecular weight genomic DNA purification kit according to the manufacturer’s protocol (QIAGEN Genomic tip either 20/G or 100/G) and Quant-iT Picogreen dsDNA quantification. Following genomic DNA isolation, the purity and quality was assessed using a Nanodrop (ND-1000). DNA damage in the mitochondrial genome was measured utilizing the Mito DNA_DX_ the assay to calculate mitochondrial DNA lesion frequency as previously described^42^. Reaction mixtures used KAPA Long Range HotStart DNA Polymerase (KAPABiosystems) in a 96-well platform. Primers used for human short and long amplicons can be found in^69^. Each biological DNA sample was performed in triplicate on two independent days (for a total of 6 PCR reactions).

### Statistical analyses

For gene expression, data was analyzed by one-way ANOVA with Dunnet’s post-test for multiple comparisons, using GraphPad software (GraphPad, La Jolla, CA). p < 0.05 was considered statistically significant.

#### Quantification of functional fluorescence signal

Raw image data from CellROX assay experiments were imported into CellProfiler for quantification.^70^ Briefly, the nuclei DNA stain (Hoechst 33342, blue channel) was used to identify individual cells. Fluorescence intensity of CellROX green stain was measured in each individual nuclei and plotted in graphs.

## Acknowledgements

We are indebted to the patient and his family for participating and donating skin samples for this study. The study was supported by awards from the California Institute of Regenerative Medicine (DISC2-09610 B.S., EDUC2-08394 B.S., D.P.), NINDS R01NS119528 (L.H.S.) and the Michael J. Fox Foundation (grant number: 16138).

## Author contributions statement

D.S., F.Z., C.A.M.T., L.H.S. and D.P. designed and conducted experiments, analyzed raw data, wrote sections of the manuscript, and prepared figures. M.R., S.Q. shared CRISPR plasmids and supported the design of the Cas9 vectors. D.K. provided feedback on design and data analysis.

B.S. conceived and designed the study, obtained funding, and wrote manuscript.

All authors reviewed and approved the final manuscript.

D.S., F.Z., C.A.M.T., and B.S. performed a portion of the work as employees at the Parkinson’s Institute and Clinical Center.

## Additional information

The authors declare no conflict of interest.

