## Supplemental Material for "Nuclease-dead S. *aureus* Cas9 downregulates alpha-synuclein and reduces mtDNA damage and oxidative stress levels in patient-derived stem cell model of Parkinson’s disease"

*corresponding author

^ Current affiliation: SRI International, Biosciences Division, Menlo Park, 94025 CA, U.S.A

*** Corresponding author**

Birgitt Schüle, MD, Dr. med.

Stanford University School of Medicine

Department of Pathology

300 Pasteur Dr., R271/217

Stanford, CA 94305

**Supplementary Table 1: Genomic position of sgRNA and transcription start sites in the promoter region of *SNCA* gene (GRCh37/hg19).**

| **sgRNA/TSS** | **Position (GRCh37/hg19)** |
| --- | --- |
| 155F | chr4: 90756784-90756804 |
| 155R | chr4: 90757811-90757831 |
| 178R | chr4: 90757834-90757854 |
| 202F | chr4: 90757831-90757851 |
| 228R | chr4: 90757884-90757904 |
| 267R | chr4: 90757923-90757943 |
| 317R | chr4: 90757973-90757993 |
| 382R | chr4: 90758038-90758058 |
| 417F | chr4: 90757073-90757093 |
| 438F | chr4: 90758067-90758087 |
| 453R | chr4: 90758109-90758129 |
| 469F | chr4: 90758098-90758118 |
| 479R | chr4: 90758135-90758155 |
| 510F | chr4: 90757139-90757159 |
| 532R | chr4: 90759188-90759208 |
| 539F | chr4: 90757168-90757188 |
| 552R | chr4: 90757208-90757228 |
| 564R | chr4: 90759220-90759240 |
| 571F | chr4: 90758200-90758220 |
| 626R | chr4: 90757282-90757302 |
| 629F | chr4: 90757258-90757278 |
| 645F | chr4: 90759274-90759294 |
| 681R | chr4: 90757337-90757357 |
| 696F | chr4: 90757352-90757372 |
| 716R | chr4: 90757372-90757392 |
| 738R | chr4: 90757394-90757414 |
| 792F | chr4: 90759421-90759441 |
| 802F | chr4: 90758431-90758451 |
| 821F | chr4: 90758450-90758470 |
| 836F | chr4: 90758492-90758512 |
| 845F | chr4: 90759474-90759494 |
| 849R | chr4: 90759505-90759525 |
| TSS1 | chr4:90,759,446-90,759,448 |
| TSS2.1 | chr4:90,758,348-90,758,350; |
| TSS2.2 | chr4:90,758,123-90,758,125 |
| TSS3 | chr4:90,757,362-90,757,364. |

TSS = transcription start site

**Supplementary Table 2: Off-target analysis summary for sgRNAs functionally tested in SNCA-triplication iPSCs**

| **TSS** | **ID** | **Genomic Position (GRCh37/hg19)** | **Off-target analysis per number of mismatches** | | | | ***SNCA* mRNA downregulation in *SNCA*-triplication iPSC** |
| --- | --- | --- | --- | --- | --- | --- | --- |
|  |  |  | 1 | 2 | 3 | 4 |  |
| 1 | 792F | chr4: 90759421-90759441 | - | - | - | 16 (2 exonic, 5 intronic, 9 intergenic) | 25% |
| 2.2 | 382R | chr4: 90758038-90758058 | - | 2 (1 intronic, 1 intergenic) | 2 (1 intronic, 1 intergenic) | 15 (8 intronic, 7 intergenic) | 75% |
| 2.2 | 228R | chr4: 90757884-90757904 | - | 1 (intergenic) | 1 (exonic) | 7 (1 exonic, 1 intronic, 5 intergenic) | 50% |
| 3 | 510F | chr4: 90757139-90757159 | - | - | - | 6 (2 intronic, 4 intergenic) | 75% |

**Supplementary Table 3: SYBR™ green primers for the detection of SNCA gene isoforms for qPCR.**

| **Target** | **Source** | **Primer** | **Primer Sequence 5’-3’** | **Product size** | **Tm °C (Avg)** | **Tm °C (Beacon)** | **Target** | **Tm °C** |
| --- | --- | --- | --- | --- | --- | --- | --- | --- |
| *SNCA*140 | McLean et al.^1^ | Fwd | AAAACCAAGGAGGGAGTGGT | 238 | 55.3 | 55.62 | exon 3 | 55 |
|  |  | Rev | TGTCAGGATCCACAGGCATA |  |  | 55.01 | exon 5 |  |
| *SNCA*126 | Bungeroth et al.^2^ | Fwd | AAAGAGGGTGTTCTCTATGTAGTGG | 185 | 56.31 | 57.84 | exon 2-4 | 55 |
|  |  | Rev | TGTGGGGCTCCTTCTTCAT |  |  | 54.78 | exon 5 |  |
| *SNCA*112 | Bungeroth et al.^2^ | Fwd | TGTCAGGATCCACAGGCATA | 178 | 55.7 | 55.88 | exon 3 | 55 |
|  |  | Rev | ATACCCTTCCTTGCCCAACT |  |  | 55.46 | exon 4-6 |  |
| *SNCA*98 | Piper | Fwd | CTCTATGTAGTGGCTGAGAAGA | 165 | 54.7 | 53.92 | exon 2-4 | 55 |
|  |  | Rev | TGTCAGGATCCACAGGCATA |  |  | 55.48 | exon 4-6 |  |
| *GAPDH* | Piper | Fwd | CATCACCATCTTCCAGGAGC | 182 | 56.2 | 55.01 |  | 55 |
|  |  | Rev | ATGACGAACATGGGGGCATC |  |  | 57.7 |  |  |

Tm, melting temperature; Avg, average.

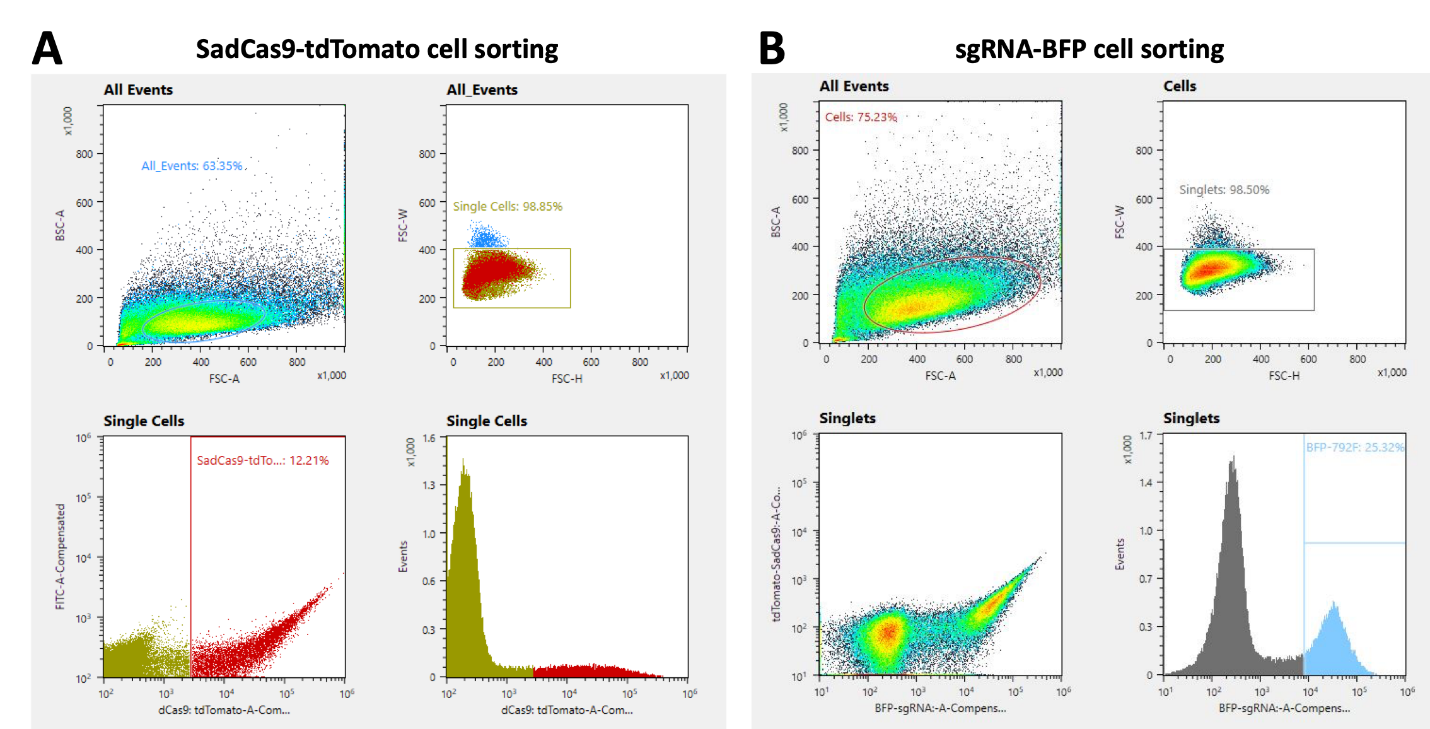

**Supplementary Figure 1:** Fluorescence-activated cell sorting of SadCas9 expression in human iPSCs. (**A**) FACS isolation of *SNCA*-triplication for SadCas9-tdTomato and (**B**) for sgRNA-BFP. iPSCs were transduced with 12.5 μL of concentrated lentivirus SadCas9-tdTomato and 12.5 μL of concentrated lentivirus rtTA in 400 μL of StemFlex medium with 6 μg/mL of polybrene. After 24 h, media was changed, and cells were cultured until they reached 100% confluency. Confluent cell cultures were expanded into 6-well plates for 1-2 weeks. To induce SadCas9 expression, cells were treated with 1 μg/mL doxycycline for 24 h prior to cell sorting. Cells were sorted based on expression of tdTomato as a marker for expression of SadCas9. Sorted cells were a heterogeneous population showing different levels of SadCas9 expression. These cells were expanded for another 1-2 weeks. To normalize the level of SadCas9 expression and allow accurate comparison between sgRNAs, SadCas9-iPSCs were clonally selected using serial dilution method. A final dilution of 0.5 cells/100 μL of medium was plated into 96-well plates. sgRNA infection of clonal SadCas9-iPSCs was performed using 25 μL of concentrated lentivirus sgRNA in 400 μL of StemFlex medium with 6 μg/mL of polybrene. Cells were expanded into 6-well plates for 1-2 weeks prior to sorting.

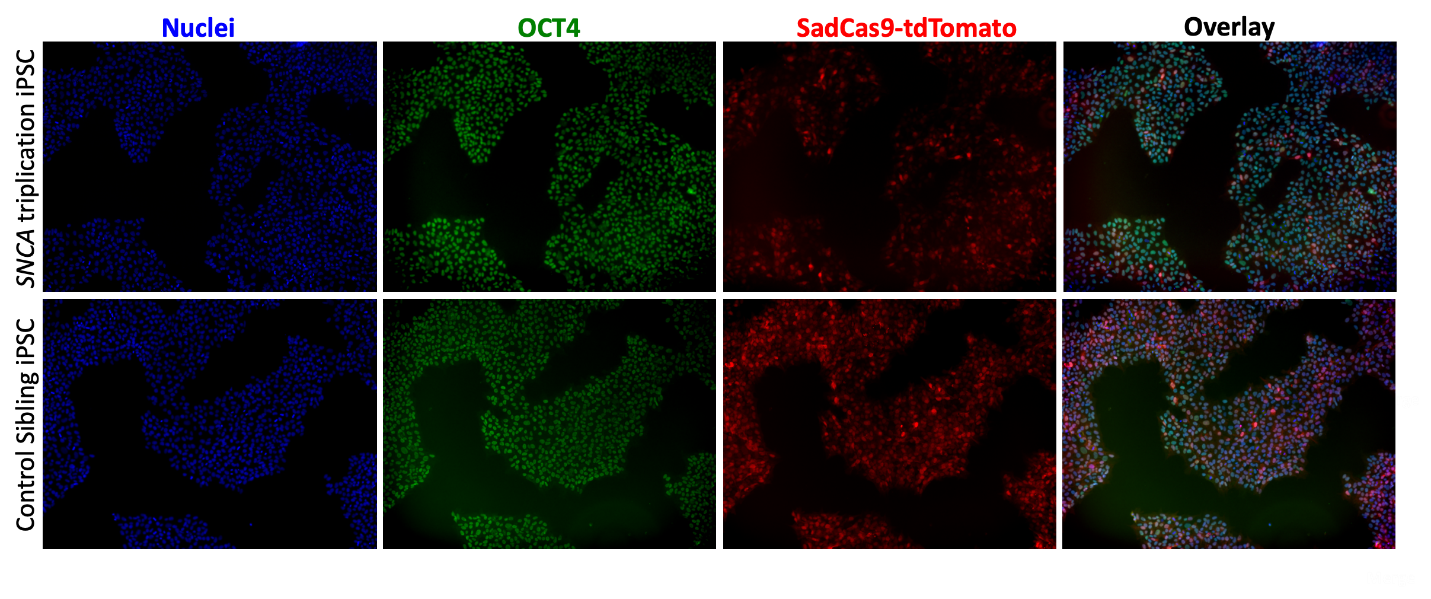

**Supplementary Figure 2:** **Patient derived iPSCs remain pluripotent after lentiviral integration of SadCas9 expression cassette.** Representative 10X images of *SNCA*-triplication iPSCs (upper panel), and control sibling iPSCs (lower panel) expressing pluripotency marker OCT4 (green) while expressing SadCas9-tdTomato (red) at 24 h post-treatment with 1 μg/mL doxycycline.

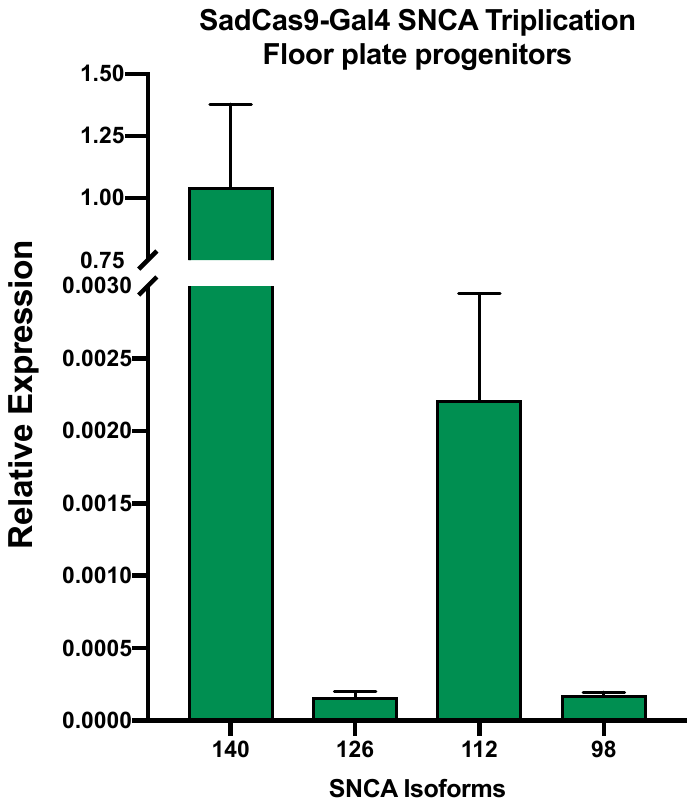

**Supplementary Figure 3. Relative expression of *SNCA* isoforms in floor-progenitor cells derived from *SNCA*-triplication iPSCs.**

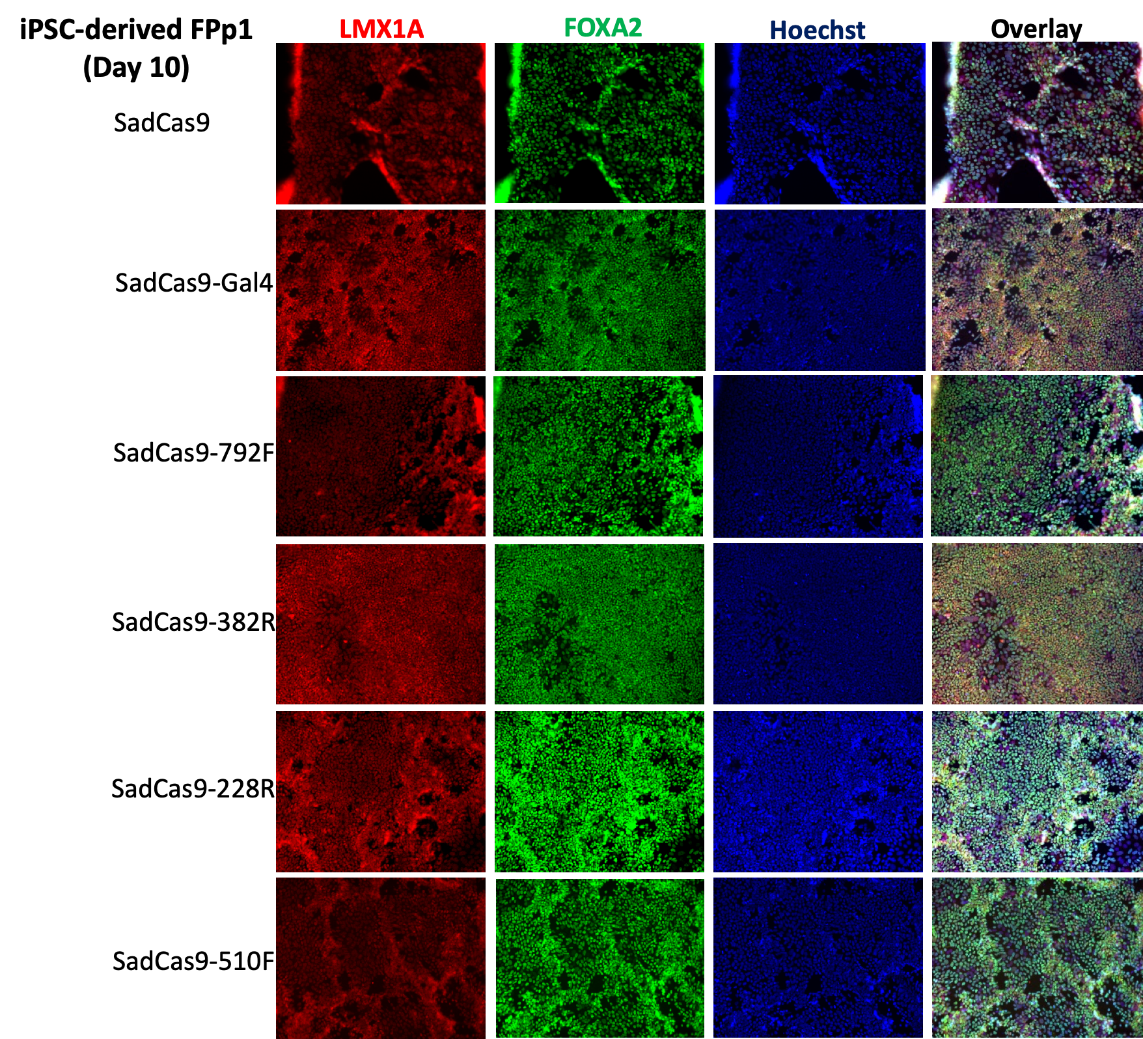

**Supplementary Figure 4. Neuronally differentiated floorplate progenitors (FPp1 at day 10 of differentiation) derived from SadCas9/sgRNA expressing iPSCs. All lines express floorplate progenitor markers LMX1A and FOXA2.**

***References***

1	McLean, J. R., Hallett, P. J., Cooper, O., Stanley, M. & Isacson, O. Transcript expression levels of full-length alpha-synuclein and its three alternatively spliced variants in Parkinson's disease brain regions and in a transgenic mouse model of alpha-synuclein overexpression. *Molecular and cellular neurosciences* **49**, 230-239, doi:10.1016/j.mcn.2011.11.006 (2012).

2	Bungeroth, M. *et al.* Differential aggregation properties of alpha-synuclein isoforms. *Neurobiol Aging* **35**, 1913-1919, doi:10.1016/j.neurobiolaging.2014.02.009 (2014).
